# Bioremediation of soils contaminated with petroleum solid wastes and drill cuttings by *Pleurotus* sp. strains under different treatment scales

**DOI:** 10.1101/588673

**Authors:** Roberto Romero-Silva, Ayixon Sánchez-Reyes, Yuletsis Díaz-Rodríguez, Ramón Alberto Batista-García, Danai Hernández-Hernández, Judith Tabullo de Robles

## Abstract

Wastes from the oil industry represent one of the sources of soil pollution with the greatest environmental impact. Both drill cuttings and crude residues are delivered to the soil and produce severe toxic effects, mainly due to the presence of polycyclic aromatic hydrocarbons. Various bioremediation technologies have been implemented in order to restore the soil quality and the natural auto depuration capabilities, amongst them: composting, bioaugmentation and biostimulation. All of these bioremediation techniques promise to be eco-friendlier and cheaper alternatives than other approaches. In this work we have evaluated several strains of *Pleurotus* sp. for their effect on the bioremediation of oil-contaminated wastes and drill cuttings disposed in storage tanks or in open-air soil lots for many years. Our results suggest that combined natural attenuation mechanism and directed fungal biodegradation activities, could be promising strategies to remediate heavily petroleum polluted soils and drilling wastes both at the laboratory and in field conditions. Furthermore, we present new data that supporting *Pleurotus* genera as able to degrade asphaltenes, the most recalcitrant fraction of petroleum. This study proposes an approach that at the same time can treat soils contaminated with waste from drill cuttings and bottoms of crude storage tanks.

## 1. Introduction

The exploitation of petroleum resources is one of the anthropogenic activities with the greatest environmental impact on aquatic and terrestrial ecosystems. Cleaning of crude oil storage tanks and drilling operations, among other activities, produce petroleum wastes that are accumulated in open-air lots, negatively impacting the soil quality. Petroleum-contaminated drill cuttings regularly contain a mix of water, oil or synthetic-based fluids with significant amounts of petroleum hydrocarbons. The hydrocarbon content includes a wide range of saturated and high-molecular-weight aromatic compounds (Broni-Bediako and Amorin 2010). Crude oil and drill cuttings are toxic, mainly due to polycyclic aromatic hydrocarbons (PAH) present in the muds (5-10%) (Petri et al. 2015). There are several physicochemical methods for drill cuttings treatment: cuttings re-injection, microwave drying, and thermal desorption. Unfortunately, all of these methods have serious technical and environmental challenges, such as expensive implementation, high energy demand, risk of accidental releases to the environment and high level of pollutant residues. Many biological processes have been investigated in order to overcome these drawbacks, e.g. composting (Ahmadi et al. 2016), bioaugmentation (Rezaei Somee et al. 2018), or biostimulation (Kogbara et al. 2016). All of these bioremediation techniques, promise to be better alternatives, eco-friendly and moneywise, than physicochemical approaches. Although bioaugmentation (the practice of adding cultured microorganisms for biodegrading soil or water contaminants), has been applied successfully in several studies of petroleum hydrocarbons biodegradation by bacteria, there are few reports of the use of processes that combine degradative activity of indigenous bacteria and the bioaugmentation with fungal specimens in soils contaminated with oil-based drill cuttings. Furthermore, fungi gather valuable metabolic characteristics as they are natural xenobiotic degraders and in fact, many species have been isolated from oil-polluted sources (Dantán et al. 2008; Marchand et al. 2017). On the other hand, they contribute with overall geomicrobial activities that play key roles in the cycling of organic matter and diagenesis (Ehrlich 2006). Their ability to be massively cultured on industrial wastes with conventional microbiological techniques, makes fungi attractive candidates for use in soil bioremediation processes.

This paper describes a comprehensive analysis in order to assess the applicability for large-scale bioremediation processes of several fungal strains, as part of the environmental management and final disposal of pollutants from the Cuban oil industry (Romero et al. 2011). The effect of four strains of *Pleurotus* sp. has been evaluated on the bioremediation of oil-contaminated soils and drill cuttings wastes disposed in storage tanks or in open-air soil lots for years, and three scales were implemented for their study at laboratory, microcosm and field levels. Our results showed that several fungal strains are capable to reduce asphaltenes and resins content in the samples tested, a task that only a few microorganisms can do.

## 2. Materials and Methods

### Fungi isolation, culture and identification

Four *Pleurotus* sp. fungal strains with abundant fruiting body, oyster-shaped and white or pale yellow in color were isolated from different decaying woods pieces collected in May 2010, from a Zoological park in Habana, Cuba. The location coordinates were 23°06’40.6″N 82°23’48.8″W. In order to obtain axenic cultures, pieces of the fruiting body were inoculated in Petri dishes with Sabouraud Dextrose Agar (BioCen Cat. 4018) and 25 μg/μL of streptomycin, at room temperature (26 ± 2°C) for seven days. Agar plugs with pure mycelium were passed to new dishes and grown for another seven days. Besides the evidence offered by the fructification bodies, the isolated strains showed morphological features (mycelia growth pattern and conidial morphology) consistent with the genus *Pleurotus*. The strains were coded as B-1, B-7, B-10, B-15 and pure cultures were conserved in the Chemistry and Biotechnology Laboratory of the Petroleum Research Center, Cuba.

### Solid State Fermentation (SEF). Laboratory scale assay

Three grams of sugarcane bagasse (particle size <1.6 mm) and 17 grams of soil contaminated with petroleum solid wastes were added to conical flasks of 100 mL capacity. They were sterilized for 15 min at 121 °C. A solution of ammonium sulfate (0.25%) sterilized independently was used to moisten each flask (32 mL). As inoculant we used two agar slants (5 mm in diameter) extracted from the active edge of colony growth of the four *Pleurotus* strains. The fermentation lasted 15 days in static condition at room temperature (26 ± 2°C). Five replicates were used for each treatment, including an abiotic control (without fungal treatment).

### Hydrocarbon Degradation in Microcosms

The microcosm assay was carried out in composters (diameter 35 cm, depth 18 cm) with sterilized petroleum solid wastes (1 kg), sugarcane bagasse (16 g) and then brought up to 60% of moisture with ammonium sulfate solution (0.25%) sterilized independently. The composters were inoculated with circular sections of mycelia (~ 20 cm^2^) from the four *Pleurotus* strains already described, with 4 fragments for each composter. The incubation period lasted 30 days at room temperature (26 ± 2°C) on static regime, with periods (6 hours) of sunlight in order to simulate field conditions. The microcosms were covered with aluminum foil and the soil was moistened by the addition of 250 mL sterile distilled water every week until the end of the experiment.

### Bioaugmentation of parcels contaminated with petroleum solid wastes and drill cuttings using a Pleurotus sp. strain

Two parcels (15×15×0.30 m) from a long-time contaminated (~10 years) region with petroleum solid wastes and drill cuttings, in La Cantera Birama, located on Matanzas province, Cuba; were selected for a field-based bioremediation experiment. First, the soil was manually conditioned by removing the surface layer. The field contained aged hydrocarbon residues and a significant rock content. Then, the parcels were biostimulated by adding sawdust according to Díaz et al. (2013) and moistened with a natural surfactant solution obtained as a byproduct of the processing of henequen fibers (*Agave fourcroydes*). Subsequently, one parcel was bioaugmented with a mixture of spores and mycelium -in Sabouraud liquid medium-from *Pleurotus* sp. B-7 strain (10E+06 spores. mL^−1^) and the other parcel was left intact, as a natural attenuation control. No physical barrier was established between the parcels. The specifications for transportation and release of the inoculum were authorized by the Biosecurity Licenses CH 17-P (78) 10 and CH17-P (81) 10 granted by the National Center for Biological Safety of Cuba (CSB). The analytical monitoring of the bioremediation process was assessed by extracting a subset of field topsoil samples (~10 grams of soil) for chemical and microbiological analysis. The following variables were recorded at 0, 30 and 70 days: total petroleum hydrocarbons (TPH) (APHA 1998), oils and grease (EPA 1999), saturated, aromatic, resins and asphaltenic components (SARA) (García-Rivero et al. 2007; Drews A 2008), total nitrogen (ISO 1995), total phosphorus (APHA 1998), total microorganism count, hydrocarbon-degrading microorganisms (Margesin et al. 2003).

### Biodegradation analysis

The rates of biodegradation expressed as a percentage were determined by monitoring the reduction of petroleum components for all samples at the initial and final stages of each experiment. Uninoculated controls supplemented with contaminated soils samples were always considered. The following equation was used for calculations:

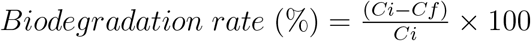

Where *Ci* indicates initial concentration; *Cf* indicates final concentration

### Statistical analysis

For each experiment a completely randomized design was carried out. The Kolmogorov-Smirnov test was used to check the normality premise. The homogeneity of variance was verified by the Chochran-Hartley and Bartlett test. The Duncan test was used for “ *a posteriori*” means comparison. The ANOVA, and two-way joining clustering analysis were assessed using the statistical package Statistica 7.0 StatSoft, Inc. (2004). Additional data are given in Supplementary material 1.

## 3. Conclusions

One of the drawbacks in bioremediation lies in achieving comparable results in laboratory as in the field. Although selected and adapted microbial strains are useful in hydrocarbon biodegradation, not all fractions are degraded in similar extension, the combination of different strategies becomes necessary in order to significantly reduce contaminant levels under realistic field conditions. Our results point toward a combined use of bioaugmentation and natural attenuation with the purpose of obtaining improved outcomes in bioremediation processes. Currently, we are carrying out studies on this site to understand how autochthonous microbial communities interact in the hydrocarbon-degradation processes, through metagenomic approaches. It is worth to highlight the isolation of a very efficient strain of *Pleurotus* capable of asphaltene degradation (B-7 strain), since very few organisms can achieve this capacity.

## 4. Results and Discussion

### Bioremediation of soils contaminated with petroleum solid wastes (Lab scale)

In order to characterize the petroleum-based hydrocarbon content in the soil, the samples were subjected to an initial analysis by standard laboratory methods for the fractionation of petroleum (Table 1). The soil samples were rich in compounds that primarily come from crude oil, being the resin, and the oils and grease, the major fractions with a content of 10^4^ mg.kg ^−1^ in average. The total petroleum hydrocarbons, saturated, aromatics and asphaltenic components oscillated in the 10^3^ mg.kg ^−1^ order. This confirmed that the soil samples were composed almost exclusively by complex mixtures of hydrocarbons derived from crude oil.

**Table 1:**
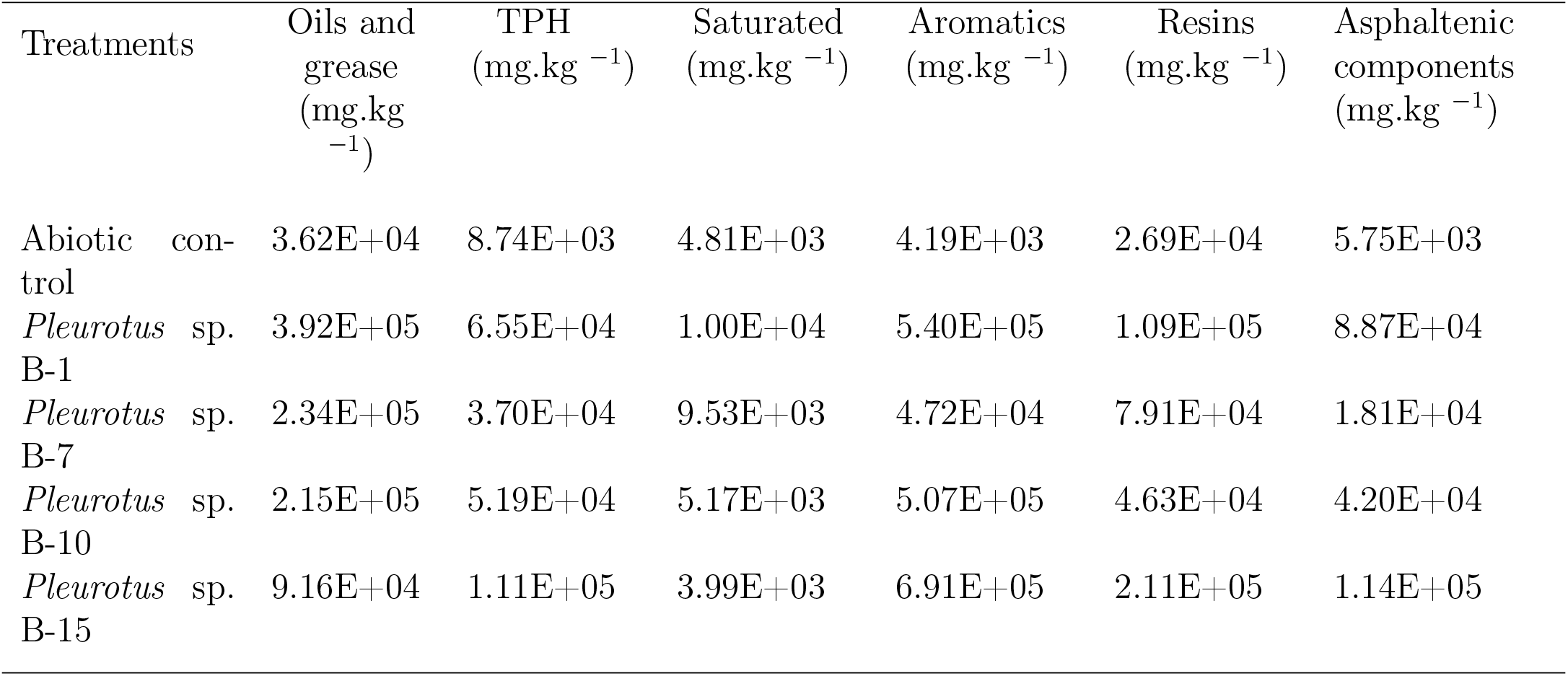
Initial characterization of soils contaminated with petroleum solid wastes

After 15 days of treatment with four strains of the basidiomycete *Pleurotus* sp. (as described in MM) we observed that all of the strains tested decreased the content of several of the oil fractions contained in the polluted soil, in contrast to the abiotic control for which no significant change was detected. The highest degradation rates were reached for saturated hydrocarbons (36%, 41.6%, 54.4% and 53.56% for the B1; B7; B10 and B15 strains, respectively) (Figure 1). The resins (maximum biodegradation of 40.5% by B10 strain), and oils and grease (maximum biodegradation of 46.34% by B15 strain) fractions were degraded by all strains. However, the aromatic fraction was highly elusive to biodegradation, with just 9.2% achieved by B-7 strain. Remarkably, we observed asphaltenes biodegradation in the experiment, being B-7 the strain with the higher biodegradation rate (46.45%) followed by B-10 strain with 21. 6%.

**Figure 1:**
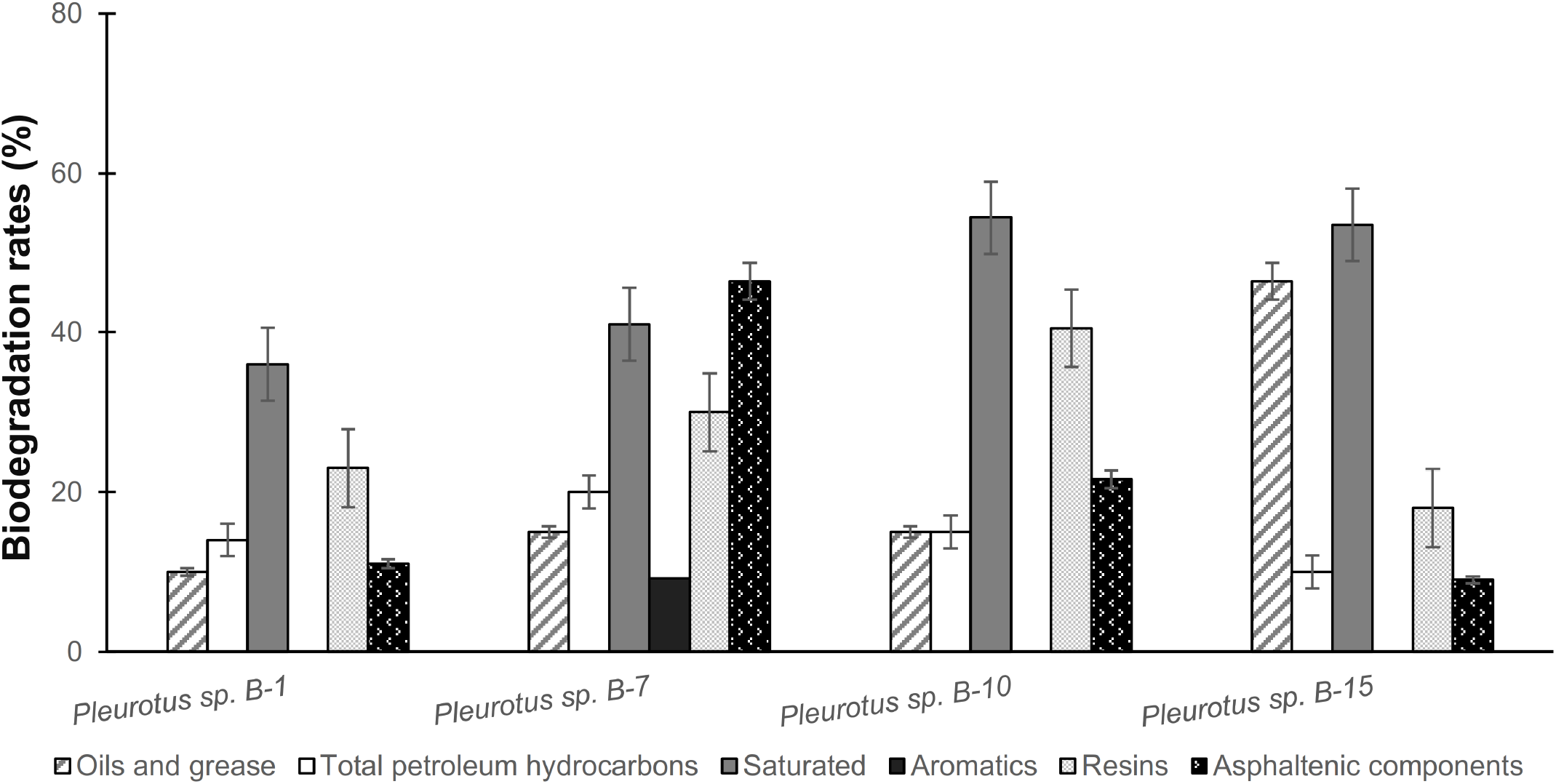
Biodegradation rates (%) for soil samples contaminated with petroleum solid wastes during 15 days of treatment with four strains of *Pleurotus* sp. (n=5)

There is very little evidence of fungi that can degrade asphalts and resins; to date, some examples have been described in detail e.g., the ascomycetes *Neosartorya fischeri* (Hernández-López et al. 2016), *Pestalotiopsis* sp. (Yanto and Tachibana 2013) and recently the basidiomycete *Daedaleopsis* sp. (Pourfakhraei et al. 2018). Asphalthenes biodegradation rates achieved by *Pleurotus* sp. B-7 and B-10 are comparable to these former examples. *Pleurotus* sp. B-7 was the only strain capable to modify all fractions in the polluted soil, a remarkable fact considering the petroleum hydrocarbons chemical complexity. For example, the resins influence the structural stability of petroleum, asphaltenes contain the highest molecular weight constituents in the crude (Andersen and Speight 2001), and aromatic compounds are very stable and resistant to biological degradation; all of these factors play a major role preventing biodegradation processes by microorganisms therefore, finding microbes capable to degradation of these fractions is valuable for the development of effective bioremediation strategies. These findings are not uncommon, for several studies support the role of *Pleurotus* for bioremediation of petroleum hydrocarbon contaminants in soil (Harms et al. 2011; Sukor et al. 2012; Mohammadi et al. 2017). The degradative activity is often associated with the ability of fungi to produce extracellular hydrolases and oxidoreductases (Varjani and Upasani 2017). We conclude that *Pleurotus* sp. B7 is a good candidate for deeper studies in bioremediation process.

### Microcosm bioremediation of soils contaminated with petroleum solid wastes

In order to scale up the laboratory soil biodegradation experiment, we carried out a microcosm soil biodegradation study for 30 days. The treatments assessed were: a non-sterile natural attenuation control (to supervise natural ability for intrinsic soil remediation in biostimulation conditions); *Pleurotus* sp. strains in contaminated soil as described in MM; and appropriate sterilized controls to assess abiotic degradation. The natural attenuation treatment achieved significant biodegradation for all fractions in the polluted soil (maximum biodegradation for saturated hydrocarbons 30.6%), with the exception of asphaltenes for which no biodegradation was detected, reinforcing the fact that native microorganisms are rarely capable of asphaltene degradation (Figure 2). Biodegradation by natural attenuation is not unusual, due to the presence of indigenous degrader microorganisms in soil samples with potential effect in intrinsic bioremediation (Lee et al. 2018). However, an effect of biostimulation should not be discarded since in real field conditions the nutritional status can be that of oligotrophy.

**Figure 2:**
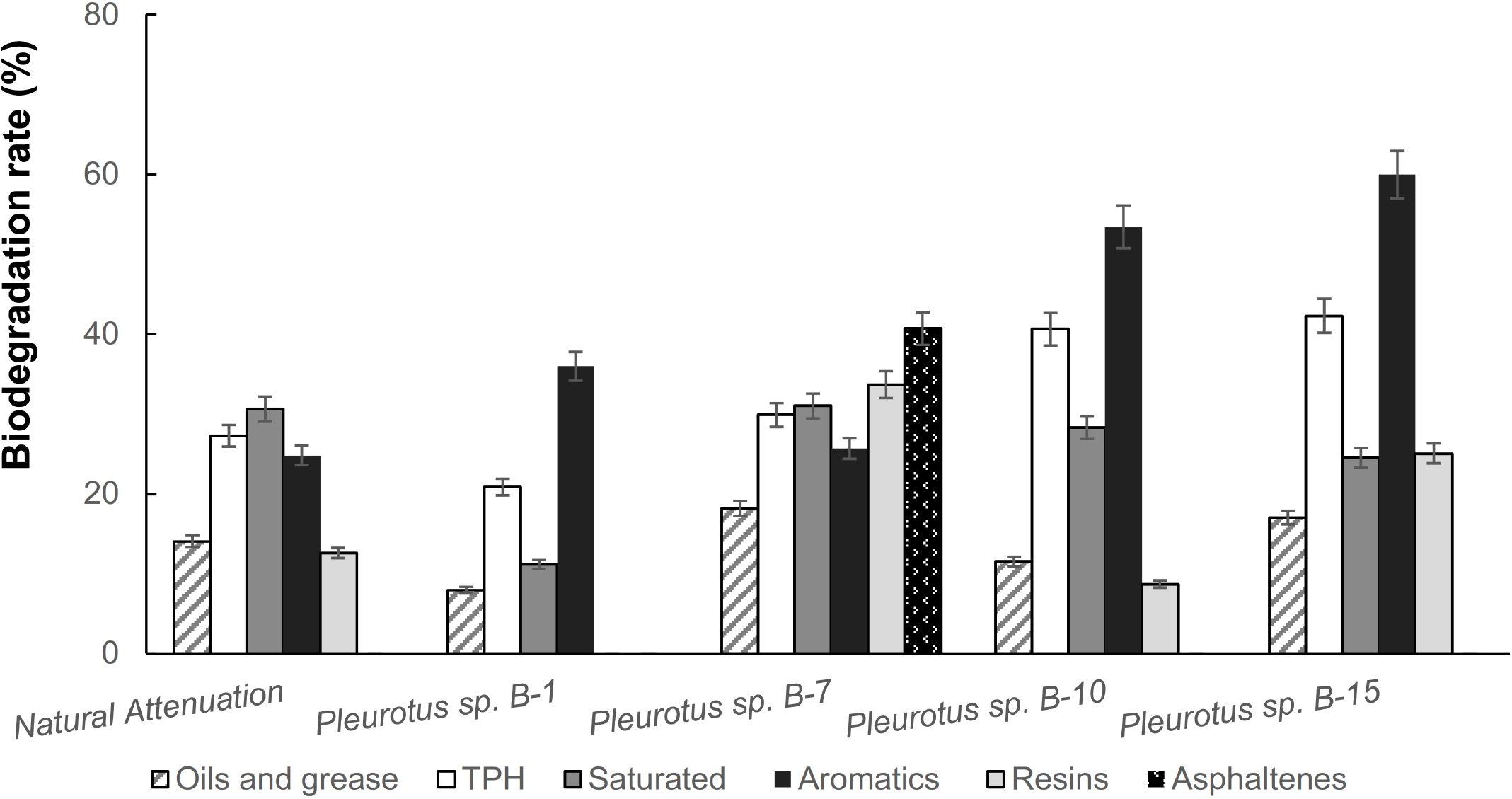
Biodegradation rates (%) achieved by four *Pleurotus* sp. strains and a natural attenuation control in soil microcosm bioremediation assay for 30 days. (n=2)

*Pleurotus* sp. B-1 could not degrade asphalts nor resins in the microcosm in contrast to our laboratory experiment, which showed a modest behavior at this respect (11-23%). *Pleurotus* sp. B-7 and B-15 degrade resins fraction considerably more than any other treatment, and the aromatics compounds were degraded by all fungal strains, contrasting with the laboratory experiment. *Pleurotus* sp. B-10 and B-15 degrade aromatic fractions in similar extension ~ 53.4-59.9% and significantly more than any other treatment. Maybe the aromatics fractions become more biodegradable when longer degradation times are applied to the process. *Pleurotus* sp. B-7 was the only one that degraded all fractions, even asphalts (40.6%), which are elusive to degradation by other treatments. The concentration of contaminants for the abiotic control by the end of the assessment showed no significant differences with the initial concentrations (in average 4.2E+04 mg.kg^−1^ vs 3.9E+04 mg.kg^−1^; standard error ~ 0.06%), suggesting that abiotic mechanisms like volatilization or dilution may not significantly contribute to degradation in these assay. These observations indicated that fungal treatment improved the biodegradation of recalcitrant fractions like aromatics, resins and asphaltenes compared to natural attenuation in the polluted soil. Furthermore, B-7 strain excels once again as a compelling candidate to perform escalated soil bioremediation studies in order to overcome natural attenuation shortcomings.

### Bioaugmentation of a field soil contaminated with petroleum solid wastes and drill cuttings with Pleurotus sp. B-7 strain

*Pleurotus* sp. B-7 was selected as study model for a real field scale bioaugmentation experiment, since it was the strain with better biodegradation performance in both the laboratory and in the microcosm experiments. Before the bioaugmentation treatment, the soil was conditioned as described in MM; a total of 10 L of the fungal inoculum was spread on the polluted soil surface. The selected soils had relatively high concentrations of petroleum solid wastes. At the beginning of experiment, the contaminant levels and hydrocarbon degrading microorganism count were quite similar for control and treated parcel according to F test (p-value > 0.05), suggesting a similar potential for natural attenuation in both sites (Figure 3A).

**Figure 3:**
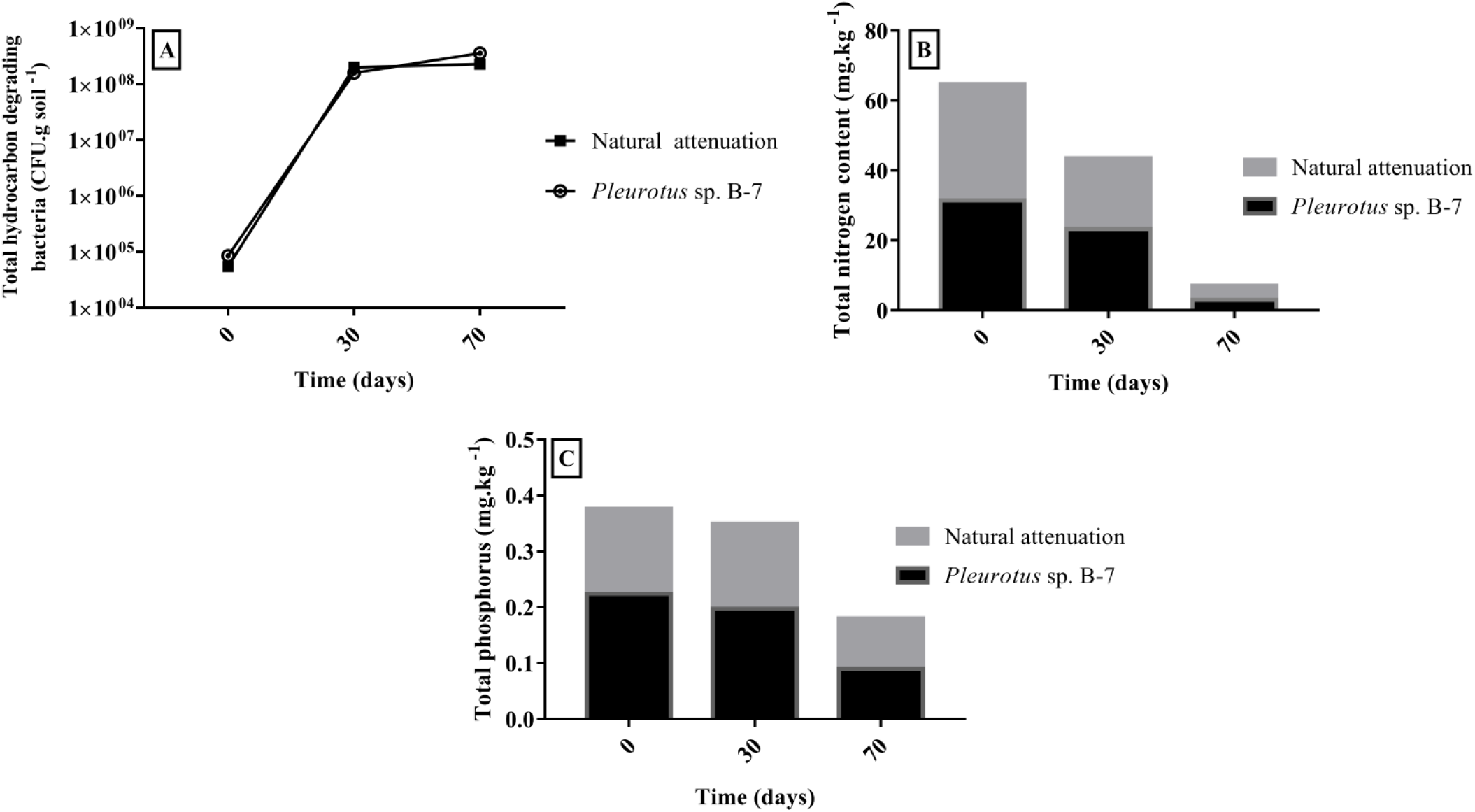
Microbial count, nitrogen and phosphorus levels in natural attenuation control and B-7 treated parcels. Total hydrocarbon-degrading bacteria (A), Total nitrogen (B), Total Phosphorus (C).

Regardless of the treatment, the main effects were observed after 70 days in terms of reducing pollutants. This was confirmed by the presence of only two statistically-different homogeneous groups (days 0 and 70) (Figure 4), clustered respect to the temporal dynamics of degradation [Oils and grease Duncan test P-value: 0.00; TPH Duncan test P-value: 0.01; Saturated hydrocarbons Duncan test P-value: 0.03); Resins Duncan test P-value: 0.01; Asphaltenic components Duncan test P-value: 0.01 (*α*: 0.05)] Supplementary material 1. Interestingly, at 70 days of treatment the Resins, Saturated and Aromatics-hydrocarbons reached accepted levels according to National Standardization Body of the Republic of Cuba (NC 819: 2017; NC 1263: 2018) (≤10000 mg. kg ^−1^).

**Figure 4:**
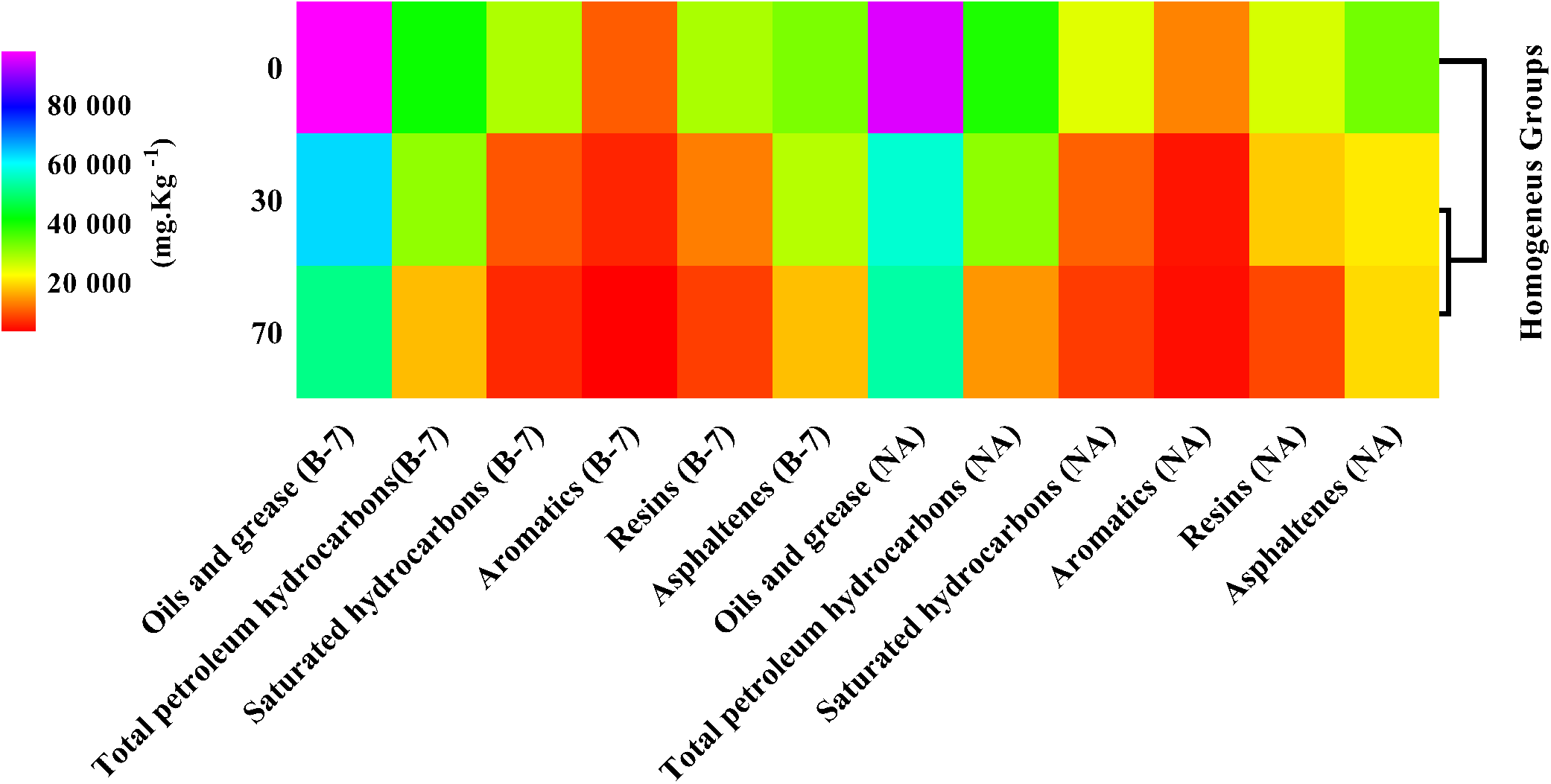
Two-Way joining graph of the oil-fractions remotion (mg.kg ^−1^) in the field assay for the natural attenuation control and B-7 strain during 0, 30 and 70 days of treatment (left side on the heatmap). Significant temporal differences between treatments are clustered in the right side dendrogram homogeneous groups (See text for P-values details and Supplementary material 1)

However, this period of time was not enough to reduce Oils and grease, or Asphaltenes up to standards accepted levels. Probably the high initial content of Oils and grease in treated parcels, interfered drastically with the microbial metabolic processes due to its diverse and complex chemical composition (coming from aged fatty matter, sulfur compounds, and organic pigments) (Pawlak and Rauckyte 2008). The percentage of resins removal achieved by the inoculum and the control, amount to 56% and 27% respectively, while the asphaltenes amounted to degrade a maximum of 14% in B-7 treatment. These findings suggest a greater resins and asphaltenes biodegradation in the parcel with the fungal inoculum, and confirm the difficulties in degrading the asphaltenics components under our field conditions. Thus, longer treatments may be necessary for accomplish the oil-pollutants levels in impacted soils, required by the normativity.

Except for TPH, resins and asphaltenes, at 70 days there were no differences between B-7 and attenuation control. The percentage of resins removal achieved by the inoculum and the control, amount to 56% and 27% respectively, while the asphaltenes amounted to degrade a maximum of 14% in B-7 treatment. These findings suggest a greater resins and asphaltenes biodegradation in the parcel with the fungal inoculum, although the contribution of indigenous microbiota is not ruled out, for example, there was a significant increase in the biodegraders count with the B-7 treatment (8.50E+04 CFU.g soil ^−1^ at 0 day and 3.55E+08 CFU.g soil ^−1^ at 70 day) (Figure 3A). This increase -in four magnitude orders-may be related with nutrient biostimulation applied to the treated parcel.

In the parcel’s topsoil, the total phosphorus concentrations were extremely low (attenuation control 0.152 mg.kg^−1^, B-7 treatment 0.224 mg.kg^−1^) at the beginning of the experiment (Figure 3C), maybe due to the historical characteristics of the field, intrinsically dry, geologically young, little fertile with sparse vegetation mostly composed by Gramineae, spiny crawling plants like *Ricinus communis*. After 70 days we noticed a decrease of phosphorous, to almost undetectable levels (~0.09 mg.kg^−1^). The total nitrogen content in both plots also suffered a significant decrease over 70 days (Figure 3B), maybe as a response to the increased metabolic demands of autochthonous and aloctone populations. Phosphorus and nitrogen are essential nutrients required by microorganism in the synthesis of cellular structures and for maintenance of adequate metabolic equilibria. The low levels detected in this study may be caused by a weak ability to release parent material rock-derived, like phosphorus, or by sequestration effects mediated by hydrophobic soil matrix contaminants (Eschenbach et al. 2013). This suggest that biostimulation plays an important role in bioremediation so maybe it should be considered for treatments at field level by adding nutrients every 70 days (in the case of this study).

There are several reports suggesting *Pleurotus* genus can effectively degrade petroleum hydrocarbons in the presence of diverse indigenous microflora (Hestbjerg et al. 2003; Mohammadi, et al. 2017). In this study, we evaluated the behavior of four strains of *Pleurotus* sp. on the degradation of polluted soils with petroleum hydrocarbons and drill cuttings. Our experimental approximation contemplated several scales to select the strain with the best performance in the bioremediation of contaminated soil. Finally, the selected strain (B-7) was tested in the field against a natural attenuation control in order to evaluate its possible use on an industrial scale to the bioremediation of soils impacted with wastes from the petroleum industry. Our strictly controlled laboratory and microcosm studies yielded promising results for biodegradation of the most of the hydrocarbon fractions in crude oil and waste. Furthermore, the field study achieved significant decrease of certain contaminants to acceptable levels at 70 days, according to the Cuban standards for crude oil residues and water-based drilling cuttings treatment (<10, 000 mg.kg ^−1^: NC 819:2017 and NC 1263: 2018). As other studies have pointed out, the combination of several bioremediation strategies could improve the hydrocarbon removal (Adams et al. 2015; Rodriguez-Campos et al. 2018). Remarkably, asphaltenic components in the polluted soils, were consistently modified by *Pleurotus* B-7 in the different experiments performed, supporting the possibility to implement an asphalt bioremediation scheme with this strain. However, these kinds of realistic field approaches require extensive environmental monitoring to weigh the real effectiveness of the soil bioremediation process in order to meet international standards. This study is the first large-scale approach to clean up soils contaminated with petroleum hydrocarbons through combined natural attenuation and bioaugmentation in the Cuban oil industry, and confirms that long term monitoring is a necessary condition in the current efforts for bioremediation of petroleum hydrocarbons polluted soils.

## Acknowledgments

We thank to Unión Cuba-Petróleo (CUPET) for the financial support; and to Jorge Luis Folch-Mallol (PhD) for his helpful comments.

