## Supplementary Material for "Bioremediation of soils contaminated with petroleum solid wastes and drill cuttings by *Pleurotus* sp. strains under different treatment scales"

[1] Centro de Investigación del Petróleo, Churruca # 481, Cerro, La Habana, Cuba.

[2] Cátedra Conacyt, Instituto de Biotecnología, UNAM, Av. Universidad 2001, Cuernavaca, Morelos, México.
[3] Centro de Investigación en Dinámica Celular, Universidad Autónoma del Estado de Morelos, México.
4 Centro de Investigación en Biotecnología, Universidad Autónoma del Estado de Morelos, México.

*To whom correspondence should be addressed

**Kolmogorov-Smirnov (K-S) test for Normality in the field scale experiment, based on the maximum difference between the sample cumulative distribution and the hypothesized cumulative distribution**

| | *N* | *max D* | *K-S (P-value)* |
|---|---|---|---|
| ***Oils and Grease*** | 12 | 0.303037 | p < .20 |
| ***TPH*** | 12 | 0.137061 | p > .20 |
| ***Saturated hydrocarbons*** | 12 | 0.224890 | p > .20 |
| ***Aromatic hydrocarbons*** | 12 | 0.213625 | p > .20 |
| ***Resins*** | 12 | 0.155288 | p > .20 |
| ***Asphaltenic components*** | 12 | 0.200822 | p > .20 |

**Tests of homogeneity of variances in the field scale experiment. Estimated effects over treatments (Natural attenuation control, *Pleurotus* B-7) at 0, 30 and 70 days**

| | **Hartley** | **Cochran** | **Bartlett** | **df** | ***P-value*** |
|---|---|---|---|---|---|
| ***Oils and Grease*** | 70.40 | 0.531185 | 3.94174 | 5 | 0.557834 |
| ***TPH*** | 205.26 | 0.571696 | 5.02313 | 5 | 0.413064 |
| ***Saturated hydrocarbons*** | 1498.44 | 0.534619 | 8.23286 | 5 | 0.143861 |
| ***Aromatic hydrocarbons*** | 459.74 | 0.426010 | 4.81066 | 5 | 0.439421 |
| ***Resins*** | 389.11 | 0.733695 | 9.42929 | 5 | 0.093119 |
| ***Asphaltenic components*** | 19777.83 | 0.977615 | 16.92699 | 5 | 0.004640 |

**ANOVA univariate results for the field scale experiment with *Pleurotus* sp. BP-7 and the Natural attenuation control. The response-variables attributes are presented. P-value < 0.05 indicate statistical significance**

| | *Sum of Squares* | *Medium Square* | *F* | *P-value* |
|---|---|---|---|---|
| | | **Oils and Grease** | | |
| *Intercept* | 5.78E+10 | 5.78E+10 | 2260 | 0.00 |
| *Treatments* | 4.55E+09 | 9.10E+08 | 36 | 0.00 |
| *Error* | 1.53E+08 | 2.56E+07 | | |
| *Total* | 4.70E+09 | | | |
| | | **TPH** | | |
| *Intercept* | 9.82E+09 | 9.82E+09 | 394 | 0.00 |
| *Treatments* | 1.17E+09 | 2.33E+08 | 9 | 0.01 |
| *Error* | 1.49E+08 | 2.49E+07 | | |
| *Total* | 1.32E+09 | | | |
| | | **Saturated hydrocarbons** | | |
| *Intercept* | 2.50E+09 | 2.50E+09 | 77.3 | 0.00 |
| *Treatments* | 8.79E+08 | 1.76E+08 | 5.4 | 0.03 |
| *Error* | 1.94E+08 | 3.23E+07 | | |
| *Total* | 1.07E+09 | | | |
| | | **Aromatic hydrocarbons** | | |
| *Intercept* | 626638721 | 626638721 | 70.6 | 0.00 |
| *Treatments* | 175659847 | 35131969 | 4.0 | 0.06 |
| *Error* | 53281902 | 8880317 | | |
| *Total* | 228941749 | | | |
| | | **Resins** | | |
| *Intercept* | 3.41E+09 | 3.41E+09 | 244 | 0.00 |
| *Treatments* | 7.45E+08 | 1.49E+08 | 11 | 0.01 |
| *Error* | 8.38E+07 | 1.40E+07 | | |
| *Total* | 8.29E+08 | | | |
| | | **Asphaltenic components** | | |
| *Intercept* | 7.55E+09 | 7.55E+09 | 926 | 0.00 |
| *Treatments* | 4.66E+08 | 9.33E+07 | 11 | 0.01 |
| *Error* | 4.89E+07 | 8.15E+06 | | |
| *Total* | 5.15E+08 | | | |

***Degree of freedom**: Intercept 1; Treatments 5; Error 6; Total 11.*

**Cumulative frequencies for response-variables (series on the right side). The bars show oils-fraction concentration ranges (mg.kg $^{-1}$) in the field scale experiment. Four-pointed stars indicate values *in range*, according to National Standardization Body of the Republic of Cuba (NC 819: 2017; NC 1263: 2018).**

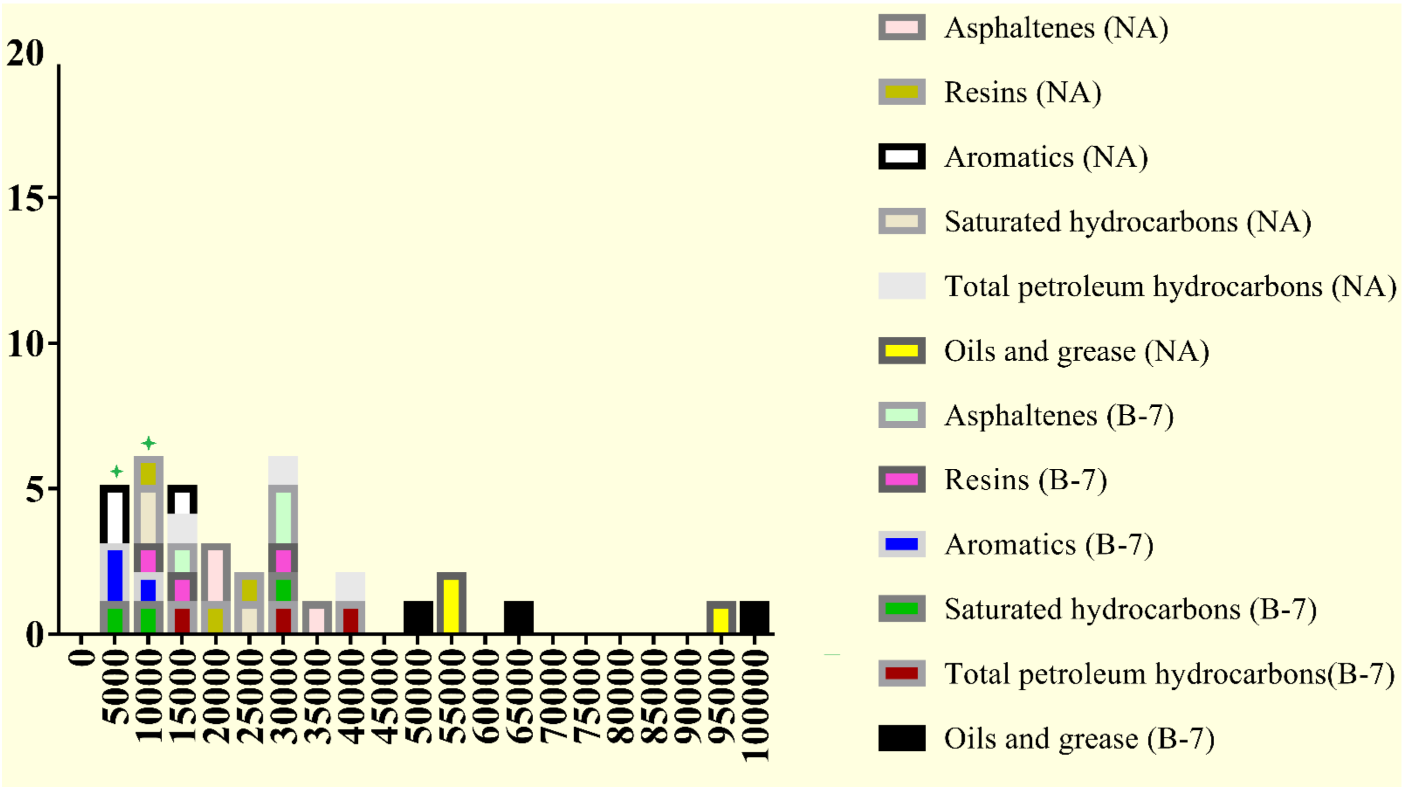
